# No major flaws in “Identification of individuals by trait prediction using whole-genome sequencing data”

**DOI:** 10.1101/187542

**Authors:** Christoph Lippert, Riccardo Sabatini, M. Cyrus Maher, Eun Yong Kang, Seunghak Lee, Okan Arikan, Alena Harley, Axel Bernal, Peter Garst, Victor Lavrenko, Ken Yocum, Theodore M. Wong, Mingfu Zhu, Wen-Yun Yang, Chris Chang, Barry Hicks, Smriti Ramakrishnan, Haibao Tang, Chao Xie, Suzanne Brewerton, Yaron Turpaz, Amalio Telenti, Rhonda K. Roby, Franz Och, J. Craig Venter

## Abstract

In a recently published PNAS article, we studied the identifiability of genomic samples using machine learning methods [Lippert et al., 2017]. In a response, Erlich [2017] argued that our work contained major flaws. The main technical critique of Erlich [2017] builds on a simulation experiment that shows that our proposed algorithm, which uses only a genomic sample for identification, performed no better than a strategy that uses demographic variables. Below, we show why this comparison is misleading and provide a detailed discussion of the key critical points in our analyses that have been brought up in Erlich [2017] and in the media. Further, not only faces may be derived from DNA, but a wide range of phenotypes and demographic variables. In this light, the main contribution of Lippert et al. [2017] is an algorithm that identifies genomes of individuals by combining multiple DNA-based predictive models for a myriad of traits.

## A perfect oracle

Erlich [2017] sets out to find a simple baseline for the problem of identification of genomic samples treated in Lippert et al. [2017]. His strategy was to categorize subjects into one of 6 ethnicities by 2 genders by 10 bins for age in increments of 7 years, *i.e.*, a total of 120 groups. Identity of an individual is predicted by random sampling of a single subject from the same group. Although this setting is a relevant baseline when assessing identifiability in a database that is associated with demographic information not protected by HIPAA, this approach assumes a-priori knowledge of age, ethnicity and gender for each subject. Thus, the setting is different from the one in Lippert et al. [2017], where the only available material and information is the genomic sample. In the latter setting, any other phenotypic and demographic variables, including the aforementioned age, ethnicity and gender, are not assumed to be present, but instead are to be predicted from the genomic sample. In the Lippert et al. [2017] setting, the Erlich [2017] result could be seen as an upper bound, that is, the result that could be achieved in an experiment where one had an oracle that allowed perfect prediction of these traits.

In reality, predictions of any phenotypes and demographic data will have limited accuracy. So the final identification results for a model using real predictions will be less accurate than for a model which assumes that all information is provided in advance, such as the Erlich [2017] model. For example, Lippert et al. [2017] report a mean absolute error for prediction of age from a genomic sample of 8.0 years. Also, a person’s ethnicity is not a perfect representation of a person’s genetic ancestry, especially in strongly admixed populations such as African Americans or Latinos, where differences in ancestry are continuous. Even for gender, we only achieved an accuracy of 99.6% when predicting the value from the genome, as reported in Supplemental Information (SI) Appendix of Lippert et al. [2017].

The “Full” model (see Table 2) of Lippert et al. [2017] operates under true biological variability and uncertainty. It also includes noise from environmental contributions and technical imprecision in the collection of traits. Therefore, the comparison with the numbers from the Erlich [2017] oracle in their Figure 1 is not informative.

## Only gender, ethnicity and age?

A question that had been raised is if predictions are only driven by gender, ethnicity and age. Lippert et al. [2017] evaluated this in detail on demographically homogeneous populations. To do so, Lippert et al. [2017] repeated identification experiments in gender and ethnicity stratified groups for Europeans and African Americans—the two most frequent ethnicities in the cohort. The results of this analysis are shown in Table 1 and SI Appendix, Figure S31, of Lippert et al. [2017].

**Table 1:** Select and match results stratified by sex and ethnicity. Shown are different numbers of pool sizes from two to 50 within (A) African American males, (B) European males (C) African American females, (D) European females. “nan” values indicate that the size of the test sample sets were too small due to stratification. For a detailed list of models and data involved in any experiment^6^, see Table 2

|  |  | African |  |  |  |  | European |  |  |  |  |  |  |  |
| --- | --- | --- | --- | --- | --- | --- | --- | --- | --- | --- | --- | --- | --- | --- |
|  |  | 2 | 5 | 10 | 20 | 50 |  |  | 2 | 5 | 10 | 20 | 50 |  |
| Male | Full | Match | 0.82 | 0.48 | 0.29 | 0.18 | nan | Full | Match | 0.72 | 0.4 | 0.23 | 0.11 | nan |
|  |  | Select | 0.74 | 0.47 | 0.28 | 0.16 | nan |  | Select | 0.67 | 0.37 | 0.24 | 0.13 | nan |
|  | All Face + Demogr. | Match | 0.8 | 0.47 | 0.29 | 0.15 | nan | All Face + Demogr. | Match | 0.74 | 0.41 | 0.2 | 0.12 | nan |
|  |  | Select | 0.75 | 0.42 | 0.27 | 0.17 | nan |  | Select | 0.7 | 0.38 | 0.24 | 0.13 | nan |
|  | All Face + Add'l | Match | 0.74 | 0.47 | 0.28 | 0.17 | nan | All Face + Add'l | Match | 0.69 | 0.35 | 0.2 | 0.088 | nan |
|  |  | Select | 0.7 | 0.44 | 0.27 | 0.15 | nan |  | Select | 0.68 | 0.36 | 0.2 | 0.13 | nan |
|  | All Face | Match | 0.77 | 0.47 | 0.28 | 0.16 | nan | All Face | Match | 0.7 | 0.38 | 0.19 | 0.1 | nan |
|  |  | Select | 0.71 | 0.42 | 0.28 | 0.15 | nan |  | Select | 0.68 | 0.36 | 0.2 | 0.11 | nan |
|  | 3D Face | Match | 0.76 | 0.39 | 0.22 | 0.15 | nan | 3D Face | Match | 0.58 | 0.26 | 0.14 | 0.082 | nan |
|  |  | Select | 0.67 | 0.37 | 0.23 | 0.14 | nan |  | Select | 0.56 | 0.25 | 0.13 | 0.073 | nan |
|  | Landmarks | Match | 0.69 | 0.33 | 0.17 | 0.093 | nan | Landmarks | Match | 0.61 | 0.24 | 0.13 | 0.053 | nan |
|  |  | Select | 0.64 | 0.34 | 0.18 | 0.086 | nan |  | Select | 0.57 | 0.25 | 0.12 | 0.07 | nan |
|  | Eyecolor | Match | 0.68 | 0.34 | 0.16 | 0.09 | nan | Eyecolor | Match | 0.73 | 0.32 | 0.16 | 0.085 | nan |
|  |  | Select | 0.63 | 0.34 | 0.17 | 0.082 | nan |  | Select | 0.7 | 0.34 | 0.19 | 0.11 | nan |
|  | Skincolor | Match | 0.62 | 0.32 | 0.17 | 0.098 | nan | Skincolor | Match | 0.54 | 0.22 | 0.1 | 0.053 | nan |
|  |  | Select | 0.6 | 0.3 | 0.18 | 0.096 | nan |  | Select | 0.51 | 0.19 | 0.087 | 0.032 | nan |
|  | Ethnicity | Match | 0.5 | 0.21 | 0.12 | 0.054 | nan | Ethnicity | Match | 0.55 | 0.2 | 0.086 | 0.05 | nan |
|  |  | Select | 0.5 | 0.2 | 0.1 | 0.052 | nan |  | Select | 0.5 | 0.2 | 0.1 | 0.05 | nan |
|  | Age | Match | 0.71 | 0.37 | 0.21 | 0.095 | nan | Age | Match | 0.61 | 0.29 | 0.14 | 0.085 | nan |
|  |  | Select | 0.67 | 0.37 | 0.22 | 0.098 | nan |  | Select | 0.6 | 0.28 | 0.16 | 0.085 | nan |
|  | Gender | Match | 0.4 | 0.28 | 0.12 | 0.17 | nan | Gender | Match | 0.42 | 0.26 | 0.17 | 0 | nan |
|  |  | Select | 0.38 | 0.28 | 0.17 | 0.1 | nan |  | Select | 0.36 | 0.29 | 0.2 | 0 | nan |
|  | Voice | Match | 0.67 | 0.34 | 0.17 | 0.072 | nan | Voice | Match | 0.56 | 0.23 | 0.11 | 0.047 | nan |
|  |  | Select | 0.61 | 0.32 | 0.16 | 0.082 | nan |  | Select | 0.53 | 0.22 | 0.12 | 0.065 | nan |
|  | Height/Weight/BMI | Match | 0.59 | 0.28 | 0.17 | 0.089 | nan | Height/Weight/BMI | Match | 0.56 | 0.21 | 0.098 | 0.052 | nan |
|  |  | Select | 0.57 | 0.28 | 0.17 | 0.099 | nan |  | Select | 0.52 | 0.19 | 0.092 | 0.05 | nan |
|  | Random | Match | 0.5 | 0.2 | 0.1 | 0.05 | 0.02 | Random | Match | 0.5 | 0.2 | 0.1 | 0.05 | 0.02 |
|  |  | Select | 0.5 | 0.2 | 0.1 | 0.05 | 0.02 |  | Select | 0.5 | 0.2 | 0.1 | 0.05 | 0.02 |
|  |  | A |  |  |  |  | B |  |  |  |  |  |  |  |
|  |  | 2 | 5 | 10 | 20 | 50 |  |  | 2 | 5 | 10 | 20 | 50 |  |
| Female | Full | Match | 0.8 | 0.5 | 0.32 | 0.21 | 0.11 | Full | Match | 0.8 | 0.52 | 0.31 | 0.17 | nan |
|  |  | Select | 0.72 | 0.46 | 0.31 | 0.2 | 0.099 |  | Select | 0.78 | 0.49 | 0.3 | 0.19 | nan |
|  | All Face + Demogr. | Match | 0.79 | 0.51 | 0.3 | 0.19 | 0.11 | All Face + Demogr. | Match | 0.82 | 0.51 | 0.32 | 0.17 | nan |
|  |  | Select | 0.73 | 0.46 | 0.29 | 0.18 | 0.097 |  | Select | 0.79 | 0.49 | 0.32 | 0.2 | nan |
|  | All Face + Add'l | Match | 0.77 | 0.46 | 0.29 | 0.18 | 0.085 | All Face + Add'l | Match | 0.84 | 0.52 | 0.29 | 0.17 | nan |
|  |  | Select | 0.71 | 0.43 | 0.27 | 0.18 | 0.087 |  | Select | 0.79 | 0.5 | 0.3 | 0.16 | nan |
|  | All Face | Match | 0.74 | 0.4 | 0.25 | 0.15 | 0.069 | All Face | Match | 0.85 | 0.51 | 0.29 | 0.12 | nan |
|  |  | Select | 0.69 | 0.39 | 0.24 | 0.14 | 0.077 |  | Select | 0.79 | 0.48 | 0.29 | 0.15 | nan |
|  | 3D Face | Match | 0.71 | 0.36 | 0.21 | 0.11 | 0.054 | 3D Face | Match | 0.71 | 0.34 | 0.19 | 0.097 | nan |
|  |  | Select | 0.65 | 0.34 | 0.21 | 0.11 | 0.045 |  | Select | 0.66 | 0.34 | 0.19 | 0.11 | nan |
|  | Landmarks | Match | 0.64 | 0.32 | 0.15 | 0.075 | 0.029 | Landmarks | Match | 0.6 | 0.31 | 0.15 | 0.09 | nan |
|  |  | Select | 0.6 | 0.29 | 0.16 | 0.077 | 0.027 |  | Select | 0.61 | 0.28 | 0.17 | 0.077 | nan |
|  | Eyecolor | Match | 0.68 | 0.32 | 0.18 | 0.11 | 0.054 | Eyecolor | Match | 0.77 | 0.39 | 0.21 | 0.097 | nan |
|  |  | Select | 0.63 | 0.32 | 0.18 | 0.1 | 0.052 |  | Select | 0.77 | 0.39 | 0.24 | 0.13 | nan |
|  | Skincolor | Match | 0.56 | 0.24 | 0.14 | 0.064 | 0.027 | Skincolor | Match | 0.52 | 0.29 | 0.14 | 0.092 | nan |
|  |  | Select | 0.55 | 0.23 | 0.12 | 0.044 | 0.019 |  | Select | 0.53 | 0.27 | 0.18 | 0.09 | nan |
|  | Ethnicity | Match | 0.47 | 0.2 | 0.11 | 0.053 | 0.023 | Ethnicity | Match | 0.54 | 0.2 | 0.1 | 0.065 | nan |
|  |  | Select | 0.5 | 0.2 | 0.099 | 0.048 | 0.021 |  | Select | 0.5 | 0.2 | 0.1 | 0.05 | nan |
|  | Age | Match | 0.69 | 0.36 | 0.2 | 0.11 | 0.058 | Age | Match | 0.69 | 0.33 | 0.16 | 0.075 | nan |
|  |  | Select | 0.67 | 0.37 | 0.21 | 0.12 | 0.048 |  | Select | 0.66 | 0.33 | 0.18 | 0.075 | nan |
|  | Gender | Match | 0.5 | 0.25 | 0.14 | 0.11 | 0.02 | Gender | Match | 0.36 | 0.32 | 0.1 | 0 | nan |
|  |  | Select | 0.52 | 0.23 | 0.14 | 0.075 | 0 |  | Select | 0.32 | 0.32 | 0.2 | 0 | nan |
|  | Voice | Match | 0.68 | 0.35 | 0.21 | 0.11 | 0.057 | Voice | Match | 0.67 | 0.34 | 0.19 | 0.12 | nan |
|  |  | Select | 0.64 | 0.34 | 0.19 | 0.11 | 0.047 |  | Select | 0.65 | 0.31 | 0.15 | 0.078 | nan |
|  | Height/Weight/BMI | Match | 0.57 | 0.26 | 0.12 | 0.064 | 0.031 | Height/Weight/BMI | Match | 0.61 | 0.25 | 0.15 | 0.087 | nan |
|  |  | Select | 0.54 | 0.24 | 0.12 | 0.05 | 0.022 |  | Select | 0.58 | 0.24 | 0.13 | 0.078 | nan |
|  | Random | Match | 0.5 | 0.2 | 0.1 | 0.05 | 0.02 | Random | Match | 0.5 | 0.2 | 0.1 | 0.05 | 0.02 |
|  |  | Select | 0.5 | 0.2 | 0.1 | 0.05 | 0.02 |  | Select | 0.5 | 0.2 | 0.1 | 0.05 | 0.02 |
|  |  | C |  |  |  |  | D |  |  |  |  |  |  |  |
|  |  | 2 | 5 | 10 | 20 | 50 |  |  | 2 | 5 | 10 | 20 | 50 |  |

**Table 2.**
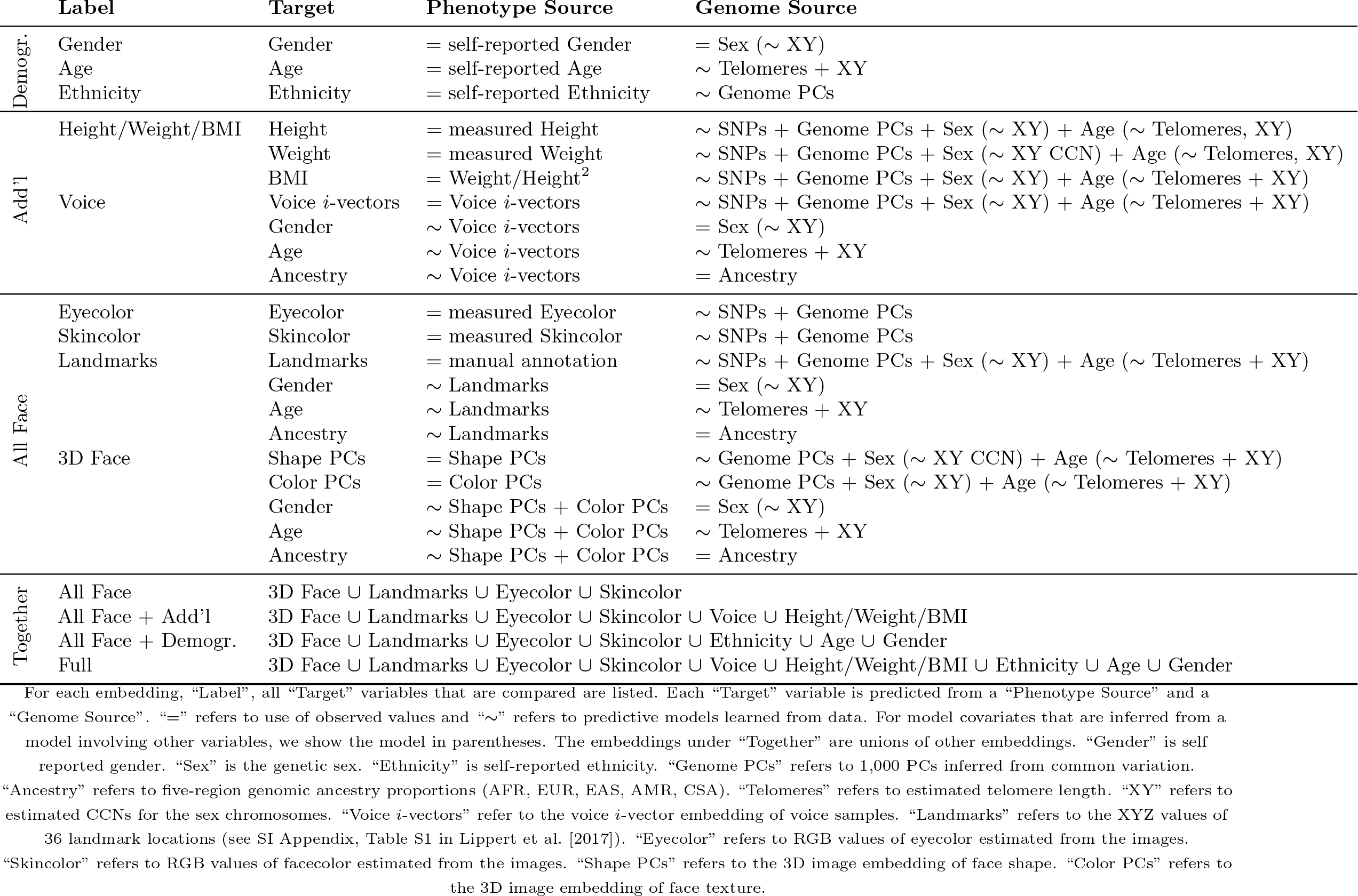
List of models used for Identification.

The only demographic variable contained in the Erlich [2017] model that is not accounted for by means of stratification is age. Within gender and ethnicity stratified pools of size 10, the “Age” model selected the correct phenotypic set out of a pool—a task we call “select” (*s*_10_)—with accuracies between 0.16 and 0.22. The model correctly matched pools of genomes to pools of phenotypic sets—a task we call “match” (*m*_10_)—with accuracies between 0.14 and 0.21. Detailed definitions of *s*_10_ and *m*_10_ are provided in Lippert et al. [2017]. These values are comparable to the “Age” model on the full cohort (see Table 2 in Lippert et al. [2017]). For the “Full” model, we have obtained *s*_10_ values between 0.24 and 0.31 in the stratified analyses, representing consistent gains between 6% and 13% over “Age”, the third variable included in the Erlich [2017] analysis. Consequently, the models provide quantifiable information beyond sex, gender and age.

## No face value?

The analysis in Lippert et al. [2017] involves a combination of many phenotypic predictors, including models derived from facial images. From facial images we derived skin color, eye color, 3D structural landmark distances, as well as 3D shape and 3D color. As images provide rich phenotypes, it is not surprising that the “3D Face” model, which consists of the shape and color components used to render faces, achieves relatively high *s*_10_ of 58% and *m*_10_ of 64% (see Table 2 in Lippert et al. [2017]). Yet, the paper does not claim that it is rendering a face from the genome that allows identification of a genomic sample, but rather a combination of many phenotypic predictors, one of which is face prediction. Indeed, when adding additional traits on top of “3D Face”, accuracy gradually increases: The “All Face” model, consisting of a combination of all traits derived from the face, increases *s*_10_ by 4% and *m*_10_ by 7%. Adding the demographic variables age, ethnicity and gender (“All Face + Demogr.”) yields additional performance increases of over 10%. This is close to the “Full” model, which also includes the remaining phenotypes not derived from facial images and has *s*_10_ of 0.74 and *m*_10_ of 0.83.

Figure S11 in the SI Appendix of Lippert et al. [2017] shows a visualization of the predicted faces for all individuals in the cohort who consented to have their images shown. For Europeans, which were genetically less heterogeneous than other ethnicities, these predicted faces are generally very similar to each other. The effect is most prominent for European males, where renderings of predicted faces look almost identical. Correspondingly, re-identification accuracies using the visualized “3D Face” variables, are consistently lower on the gender-stratified European cohorts than on the gender-stratified African American cohorts, and lowest on European males, where we obtained *s*_10_ = 0.13 and *m*_10_ = 0.14 (see Table 1B). These values are better than random, which implies that even visually similar predictions carry quantifiable information. As clearly stated in the abstract of our original work, the main contribution of Lippert et al. [2017] is not identification by face prediction, but rather an “algorithm that integrates *multiple* predictions to determine which genomic samples and phenotype measurements originate from the same person.”

## Age can be predicted at minimal cost

Erlich [2017] inaccurately points out that the approach for age prediction in Lippert et al. [2017] is expensive. His reasoning is that our approach for telomere length estimation, one component of age prediction, made use of 512 repeatedly sequenced NA12878 DNA reference samples^1^ to determine an optimal threshold to classify next generation sequencing reads as telomeric.

NA12878 sequences are a regular by-product of any large genomics facility, as the NA12878 reference DNA samples are routinely used as quality controls in sequencing [Zook et al., 2016]. Therefore, the resources used to determine this parameter, which only has to be determined a single time, are readily available. The fact that sequencing costs are going down rapidly, renders it unlikely that cost will remain a barrier, even for smaller sequencing facilities. The full result of our analysis is included in the SI Appendix, Figure S1A of Lippert et al. [2017] and may be applied and re-used in other studies. Otherwise, every sample in our study has been sequenced only once at a depth of 30X, and Ding et al. [2014] as well as SI Appendix, Figure S2 of Lippert et al. [2017] show that a minimum depth of as low as 2.5X is sufficient to determine telomere length. To conclude, telomere length prediction adds insignificant overhead to a routine whole genome sequence.

## Let’s face it!

Our work is a proof of concept done with a small set of individuals that presents a strategy for identification of genomic data using predictive modeling. While facial images provide rich data from which to build phenotypic models, it is important to notice that it is not only faces that may be derived from the genome. The genome codes for a vast number of heritable phenotypes. For many of these, larger data sets of genomic and phenotypic data will improve prediction accuracy. The genome also provides information about demographic variables, such as gender, ethnicity, and age, plus others not treated in this study, such as surnames [Gymrek et al., 2013]. Lippert et al. [2017] presents an algorithm that enables optimal combination of any number of such phenotypic and demographic prediction models. Each of these models contributes additional identifying information. Gradual improvements derived from larger data sets and better models are central to the success of machine learning applications.

In the current public debate of our work, it is useful to separate the scientific results and the importance/implications of these results. For the former, the question is whether we correctly applied and conveyed what we set out to do. Hopefully we did. For the latter, the question is whether the ideas, methods or results turn out to be important. This discussion demonstrates that it is a sensitive topic triggering a sometimes emotional debate. We definitely welcome a constructive debate on the topic of genomic privacy, a topic that is of relevance for policy and for genomic sciences: “Although sharing of genomic data is invaluable for research, our results suggest that genomes cannot be considered fully deidentifiable and should be shared by using appropriate levels of security and due diligence.” [Lippert et al., 2017] As the community relies on the availability of genomic data to decode the genetic causes of disease, we should openly face this challenge and discuss potential implications for personal privacy. Moreover, we emphasize the importance of advanced genome privacy protection strategies^2^ for enhancing data exchange.

## Footnotes

1 http://jimb.stanford.edu/giab/

2 https://genomeprivacy.org

